# A Novel Cell-Type Specific Circadian Reporter Mouse Reveals Self-Sustained Food Entrainable Nature in Enteric Neurons

**DOI:** 10.1101/2025.11.16.688705

**Authors:** Isabel Magaña, S. K. Tahajjul Taufique, Yongli Shan, Melody Shen, David E. Ehichioya, Joseph S. Takahashi, Shin Yamazaki, Yuuki Obata

## Abstract

Global luciferase reporter gene technology is an important real-time readout for the analysis of circadian oscillations. However, because nearly all cells in the body possess cell-autonomous circadian oscillators, developing cell-type–specific reporter systems is essential to dissect how these oscillators interact within complex multicellular tissues and are modulated by brain-body circadian signals. The intestine is a complex organ composed of diverse cell types with distinct origins and functions, and it exhibits robust daily rhythms in physiological activities driven by intrinsic circadian clocks. While the circadian regulation of the mucosal compartment, including the intestinal epithelium, has been relatively well characterized, the mechanisms governing the circadian rhythms in the cells of the muscularis externa remain largely unexplored. Here, we report a novel Cre-dependent Per2-luciferase reporter mouse that enables precise measurement of circadian oscillations in specific peripheral cell populations *ex vivo* and demonstrate its utility in revealing the hierarchical circadian chrono-architecture within the intestine. *Ex vivo* gut explants from mice expressing the reporter in one of five major cell types of the muscularis externa—enteric neurons (ENs), enteric glial cells (EGCs), interstitial cells of Cajal (ICCs), smooth muscle cells (SMCs), and muscularis macrophages (MMs)—exhibited robust, self-sustained circadian bioluminescence rhythms, indicating that all of these cell types possess cell-autonomous circadian oscillators. Notably, ENs in the small intestine entrained more rapidly to feeding schedules than did those in the colon, revealing regional differences in food entrainment among the gut clocks. Moreover, the clocks of ENs, SMCs, and MMs shifted their phase in response to daytime-restricted feeding, whereas ICC clocks remained unaffected even after three weeks of restricted feeding. These findings demonstrate that distinct intestinal cell types possess unique entrainment properties and that feeding rhythms can induce heterogeneous phase shifts within the gut clocks. Given that circadian disruption, such as that caused by shift work, contributes to intestinal disorders, this reporter system could provide a powerful platform to dissect the mechanisms linking intercellular circadian desynchrony to intestinal homeostasis.

**Graphical Abstract:** 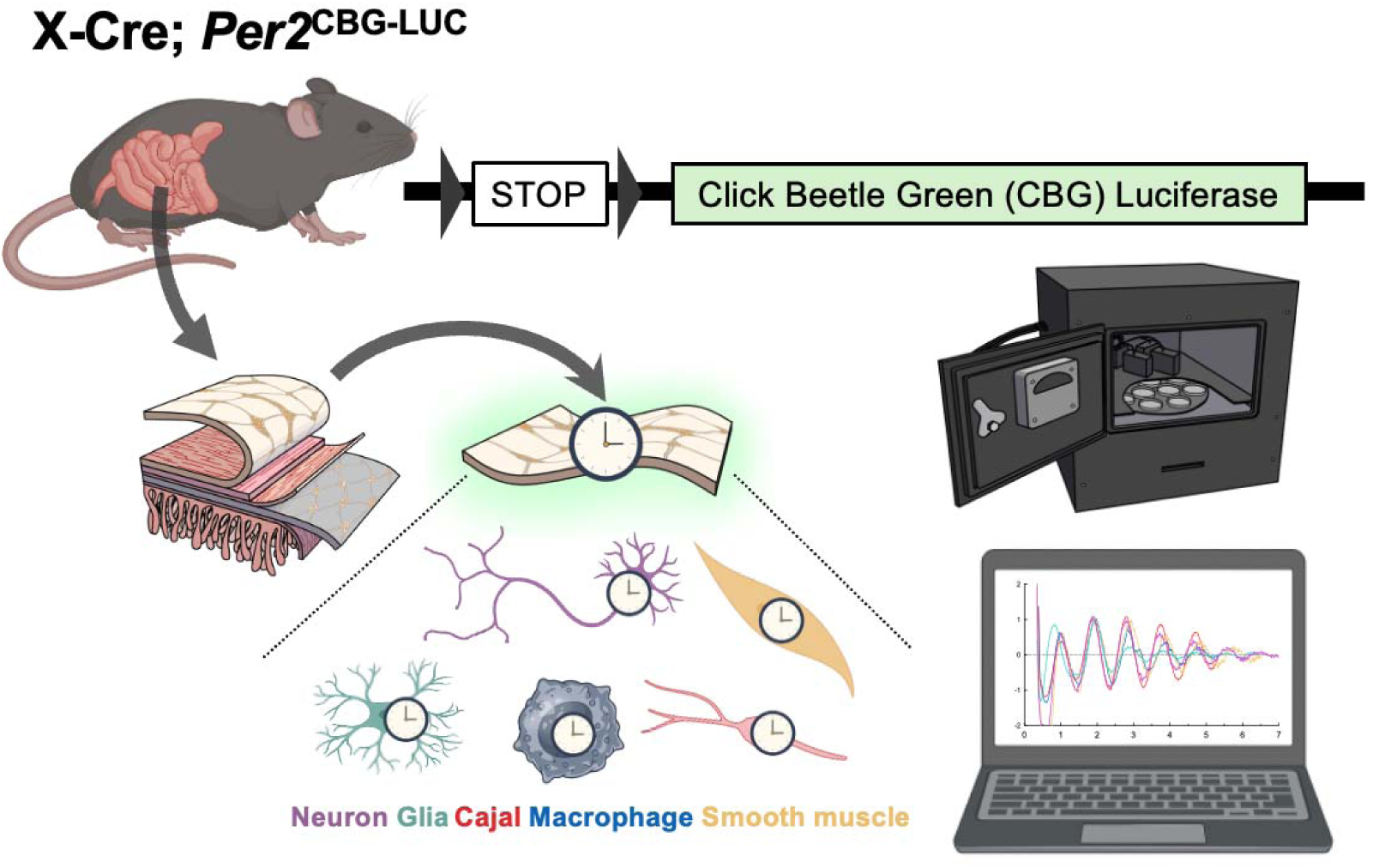

**Highlights:**

- A cell-type–specific circadian reporter mouse enables dissection of the chronoarchitecture of complex multicellular organs (e.g., the intestine)
- All five major intestinal cell types contain autonomous circadian oscillators peaking at night
- Enteric neurons in small intestine entrain to feeding cycles faster than that in colon
- Interstitial cells of Cajal in both the small intestine and colon do not entrain to feeding cycles
- Chemogenetic activation of enteric neurons mimics food entrainment, shifting their circadian phase.

## Introduction

The mammalian circadian system is an intrinsic timekeeping mechanism that governs 24-hour cycles in physiological and behavioral processes^1^. Both forward and reverse genetic approaches successfully identified essential circadian genes in mammals^2,3^ and revealed complex regulatory networks composed of positive and negative transcription-translation feedback loops that generate cell-autonomous circadian rhythms^4^. The analysis of circadian gene promoters has provided a unique opportunity to monitor circadian gene expression in real-time using reporter systems. Following the generation of the first *Period1*-luciferase transgenic rats and the subsequent PER2::LUCIFERASE translation reporter knock-in mice^5,6^, these global circadian reporter models have revealed several fundamental properties of the mammalian circadian system. These include the presence of autonomous circadian oscillators in peripheral organs and the food-entrainable nature of those in certain tissues, as well as the critical role of the suprachiasmatic nucleus (SCN) as the central pacemaker in the brain that synchronizes peripheral rhythms to anticipate daily environmental changes and optimally coordinate physiological functions^7–9^. While the SCN-driven brain-body circadian hierarchical network is well established, how individual cellular clocks within a peripheral organ synchronize to optimize its function at appropriate times remains unclear.

The intestine is exquisitely sensitive to the cycles of day and night that govern life on earth. For example, the intestinal lining is in rhythmic contact with food depending on host sleep-wake cycles, which determines the timing of nutrient absorption, digestive processes, and intestinal motility^10–12^. Additionally, as enteric pathogens may be ingested with food, the intestine must maximize immune defense during feeding, when the risk of infection is high^13^, while simultaneously activating neural and muscular responses to support digestion and motility. Conversely, the high energetic costs of these responses necessitate their downregulation when such challenges are absent, such as during sleep. Thus, the intestine must have a robust timekeeping system that anticipates changing environmental conditions and coordinates the activities of diverse intestinal cell types, ensuring that essential functions occur at the appropriate times. Notably, misalignment between the internal clock and the external day-night cycle, such as during night shift work, has been associated with various intestinal disorders^14–19^. Despite their clinical importance, it remains unclear how clocks in distinct intestinal cell types are synchronized to coordinate organ-level functions and how circadian misalignment contributes to disease development. This knowledge gap largely stems from the lack of tools that enable cell-type–specific analysis of circadian clocks within the intestine.

Here, we report a novel reporter mouse line in which click beetle green luciferase (CBG-LUC) is expressed from the mouse *Per2* locus in a Cre-dependent manner. This system enables real-time monitoring of circadian bioluminescence rhythms from specific cell types within an organ *ex vivo*. By crossing the reporter with various cell-type–specific Cre drivers, we recorded, for the first time, self-sustaining circadian rhythms of clock gene expression in five major cell types in the intestine. We observed distinct patterns of circadian phase shift among them in mice subjected to daytime-restricted feeding. This approach provides a powerful tool for dissecting the chrono-architecture of the intestine and can help address how distinct cellular clocks synchronize with environmental changes, including shifted light cycles, feeding schedules, and local physiological cues.

## Results

### Generation of Cre-dependent PER2::CBG-LUC reporter mice

To monitor circadian rhythms in specific cell types in the intestine, we generated mice in which click beetle green luciferase (CBG-LUC) is expressed as a fusion with the endogenous PER2 protein upon Cre-loxP-mediated recombination (*Per2*^CBG-LUC^ mice) (**Fig. 1A**). This system allows real-time monitoring of circadian bioluminescence rhythms in specific Cre-expressing cell types within tissue explants over time. The targeting construct used to generate these Cre-dependent CBG-luciferase reporter mice (*Per2*^CBG-LUC^) is shown in **Supplemental Figure 1**.

**Fig. 1.**
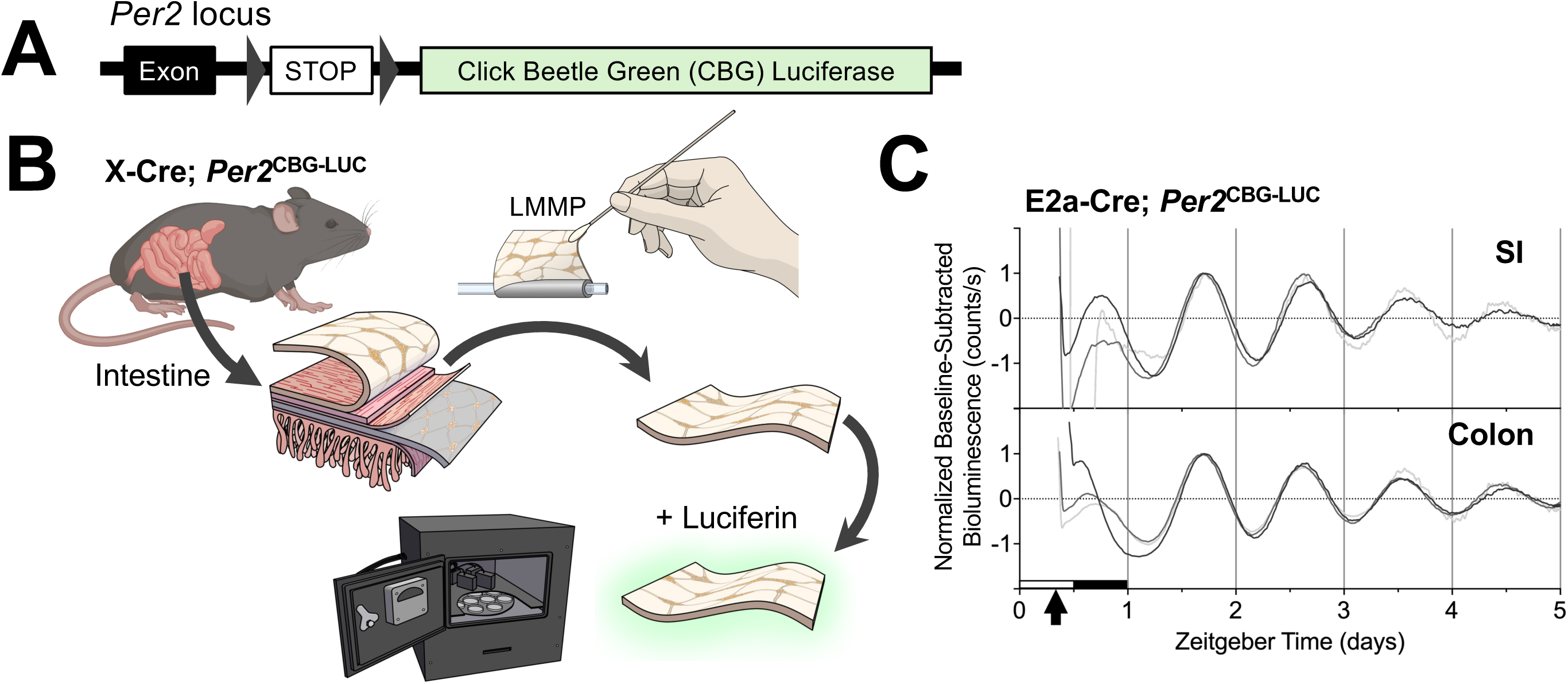
Generation of a Cre-dependent PER2::CBG-LUC reporter mouse. (A) Diagram of the PER2::CBG-LUC knock-in construct targeted to the *Per2* locus. Triangles indicate the loxP sites flanking the STOP cassette. (B) Schematic representation of the method to measure circadian rhythms from cultured longitudinal muscle–myenteric plexus (LMMP) layer of the intestine. (C) Representative normalized PER2::CBG-LUC bioluminescence traces from LMMP tissues of the small intestine (SI, top) and colon (bottom) obtained from *E2a-Cre; Per2^CBG-LUC^* mice. Baseline-subtracted bioluminescence data were normalized using the first peak after 12 h of recording as 1 (all luminescence values were divided by the value of that peak). Three representative examples per gut segment are overlaid. Black arrow represents the approximate time tissue explants were made.

The intestine is a complex organ composed of multiple layers, including epithelial, mucosal, and muscularis layers, each containing distinct cell types in varying proportions. To enable continuous *ex vivo* monitoring of circadian rhythms in intestinal cell types, we aimed to establish tissue-layer explants that preserve multiple cell populations and remain viable in culture for several days. Among these layers, we focused on the muscularis externa, which contains multiple cell types relevant to circadian physiology, including enteric neurons (ENs), enteric glial cells (EGCs), smooth muscle cells (SMCs), interstitial cells of Cajal (ICCs), and muscularis macrophages (MMs). These cell types can be dissected together in an intact preparation as the longitudinal muscle–myenteric plexus (LMMP) layer, in which the longitudinal smooth muscle and the associated myenteric plexus are preserved (**Fig. 1B**). In contrast to the rapidly renewing epithelial cells covering the mucosal surface, cells within the LMMP can be maintained for extended periods *ex vivo*^20^. Intestinal motility, driven by bidirectional interactions among ENs, ICCs, and SMCs, shows robust circadian rhythmicity^21,22^, rendering the LMMP an excellent model to study gut circadian chrono-architecture.

To validate the newly generated circadian reporter knock-in mouse, *Per2*^CBG-LUC^ mice were first crossed with the germline Cre driver E2a-Cre^23^ to generate mice globally expressing the CBG-LUC reporter. Following established methods^24^, LMMP layers were manually isolated from approximately 2 cm segments of the distal small intestine (SI) and proximal colon and were then cultured in medium containing luciferin for bioluminescence recording using a luminometer^25^ (**Fig. 1B**). When explants from these mice were cultured, robust autonomous circadian rhythms of PER2::CBG-LUC luminescence were observed in both segments, indicating the presence of circadian oscillators within LMMP cells (**Fig. 1C**). Notably, rhythms could be reliably recorded for at least one week, and, in some cases, remained stable for several weeks without changing the culture medium. These observations confirm that the newly generated *Per2*^CBG-LUC^ mice enable reliable monitoring of circadian gene expression rhythm in LMMP explants. The phase of PER2::CBG-LUC rhythm in the SI and colon was similar to those observed in explants from previously developed PER2::LUC mice (*Taufique et al, under review*), although the recording media used in these two studies are different (10% FBS vs. B27) (**Supplemental Table 1**). Of note, circadian gene expression rhythms were also detected in other peripheral organs, including the liver, adrenal gland, esophagus, kidney, and spleen, all of which exhibited phase relationships consistent with previous report^26^ (**Supplemental Table 1**), thereby validating the reliability of the novel reporter system.

### Detection of self-sustaining circadian rhythms in various intestinal cell types

Next, we sought to measure circadian bioluminescence rhythms in a cell-type–specific manner. *Per2*^CBG-LUC^ mice were crossed with cell-type–specific Cre drivers to target ENs (BAF53b-Cre)^27^, EGCs (PLP1b-CreERT2)^28^, ICCs (cKit-CreERT2)^29^, MMs (Csf1r-Cre)^30,31^, or SMCs (Myh11-CreERT2-RAD)^32^. As an alternative approach to achieve EN-specific Cre expression, we also intravenously injected *Per2*^CBG-LUC^ mice with AAV9-CaMKII-Cre, which has been shown to efficiently transduce the majority of ENs in both SI and colon^33^.

Although these Cre drivers and the AAV-based Cre delivery method have been previously validated for targeting specific cell types in the intestine, we further confirmed their specificity and efficiency in our experimental context using the R26-Sun1/sfGFP reporter line^34^. In this reporter, the nuclear envelope of Cre-expressing cells is labeled with GFP, enabling visualization of Cre activity in intestinal cell types. As expected, GFP signal was confined to the nuclei of the intended cell populations (**Supplemental Fig. 2**), confirming that our Cre strategies reliably target these cells in our experimental context.

LMMP layers isolated from each cell-type–specific PER2::CBG-LUC reporter line were cultured in medium containing luciferin (**Fig. 1B**)^25^. The cumulative total luminescence count of the five cell-type–specific reporters approximately matched the luminescence count of the global *Per2*^CBG-LUC^ reporter (**Fig. 2A**), suggesting that PER2 reporting was specific for each targeted cell population. Importantly, each cell type exhibited robust circadian rhythms that persisted for several days until recordings were terminated (**Fig. 2B**). These findings demonstrate that self-sustaining circadian oscillators exist in the five major cell types embedded within the muscularis externa of the intestine.

**Fig. 2.**
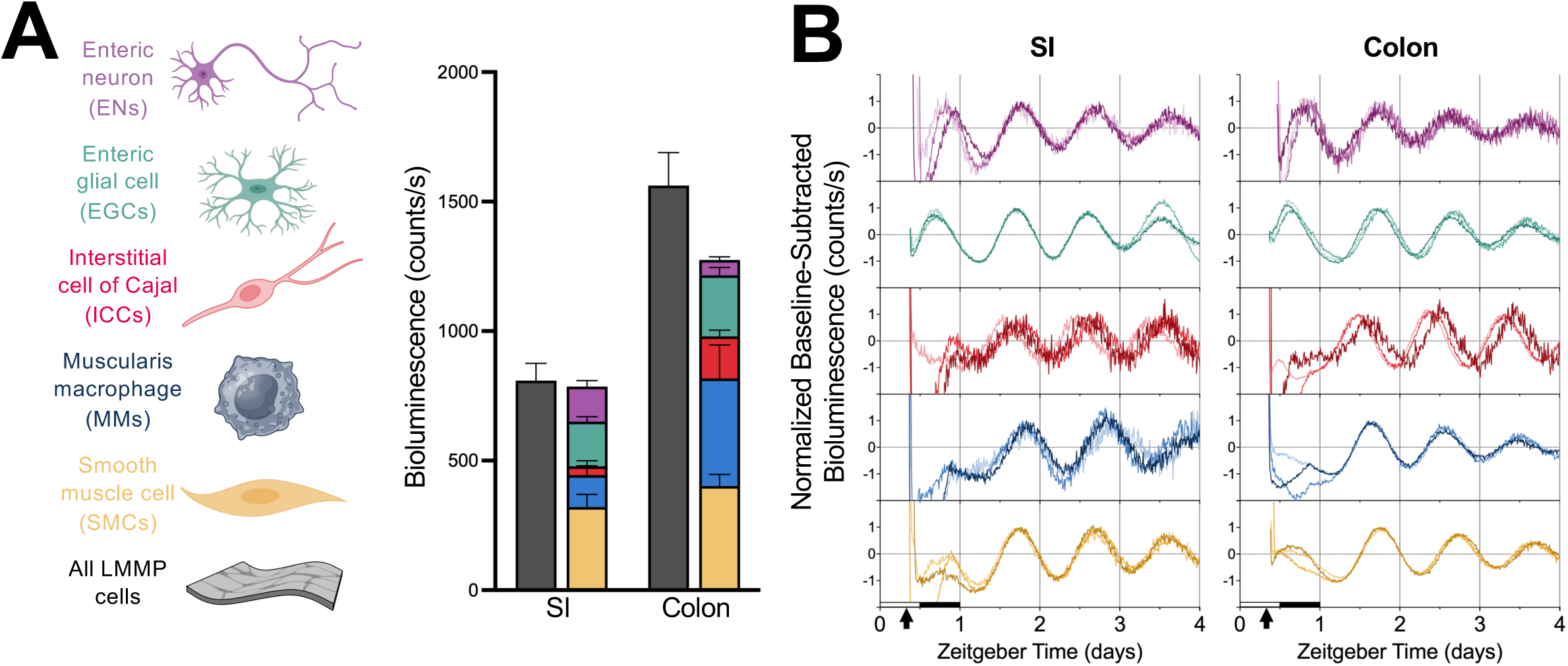
All five major cell-types in the *ex vivo* gut explants exhibited autonomous circadian rhythms. (A) Cumulative PER2::CBG-LUC bioluminescence from enteric neurons (ENs; n = 16), enteric glial cells (EGCs; n = 21), interstitial cells of Cajal (ICCs; n = 9), muscularis macrophages (MMs; n = 10), and smooth muscle cells (SMCs; n = 16), approximately matched to luminescence from the global PER2::CBG-LUC reporter (gray, n = 20) in both the small intestine (SI) and colon. Dark count–subtracted bioluminescence (counts/sec) of 2 days (hours 18–66 from ZT0 on day of explant) of recording were averaged within each sample, and then the means of all samples were averaged. (B) Representative normalized PER2 bioluminescence traces of five intestinal cell types in the SI (left) and colon (right). Baseline-subtracted bioluminescence data were normalized using the first peak after 12 h of recording as 1 (all luminescence values were divided by the value of that peak). Three representative traces obtained from PM-cultured explants for each gut segment are overlaid. Black arrows represent the approximate time when tissue explants were made.

To examine PER2 rhythms in the submucosal plexus of the ENS and associated intestinal cells, we further removed the epithelial layer from LMMP-stripped tissue, using fine needles to minimize potential effects from dead epithelial cells during culture, and performed the same recording procedure as for LMMP tissues. While PER2::CBG-LUC signals were readily detected in the global, SMC-, and MM-specific reporters, bioluminescence was not reliably observed in the EN- or EGC-specific reporters (**Supplemental Fig. 3**).This is likely due to the low abundance of these cell types in the remaining tissue, resulting in luminescence levels below the detection threshold of the photomultiplier tube. These experiments highlight the importance of efficient Cre induction and sufficient amounts of tissue for reliably detecting bioluminescence from target cell types.

### Determination of the In Vivo Phase of Circadian Clocks in Specific Intestinal Cell Types

We aimed to determine the *in vivo* phase of circadian PER2 expression in each cell type by measuring bioluminescence rhythms in cultured tissue explants. With this method, the culture procedure itself has the potential to affect the *ex vivo* phases of the explanted tissues. Estimation of *in vivo* circadian phase from *ex vivo* tissue explants relies on the assumption that these culture procedure effects are minimal and the explanted tissues retain their *in vivo* circadian phases. To test the effects of the culture procedure, LMMP tissues from neuron-specific PER2::CBG-LUC reporter were collected at two distinct time points: in the morning and evening, approximately 8 hours apart at ZT2–4 and ZT10–13 respectively (ZT0 is defined as the time of lights-on). If dissection procedures alter the phase of *ex vivo* rhythms, cultured tissues cannot be used to estimate the *in vivo* phase. Thus, this control was essential to verify that the phases measured in explanted tissues faithfully reflected the endogenous circadian states of each cell type.

When peak times were plotted relative to culture start times, most cell types showed an expected ∼8-hour difference between AM and PM dissections (**Supplemental Fig. 4**), indicating that the culture procedure did not reset these gut clocks and that *in vivo* circadian phases were preserved in LMMP explants. In contrast, a subset of colonic EGCs exhibited overlapping phases between AM and PM explants. Although the culture procedure did not completely reset the phases of these tissues, this observation suggests that colonic EGCs may be more susceptible to culture-induced phase resetting than the other cell types examined. Therefore, caution is warranted when interpreting the data from colonic EGCs.

By measuring the time of peak PER2 expression in LMMP tissues collected at both time points, we determined the phase of each intestinal cell-type–specific clock, all of which peaked during subjective night (**Fig. 3**, top plot of each cell: *ad lib*). Consistent with this, the phase of all LMMP cell types determined using the global reporter also showed a peak at subjective night (**Supplemental Fig. 5**). These findings indicate that, for most intestinal cell types, the culture procedure had minimal effect on the circadian phase, allowing us to approximate the *in vivo* phase based on *ex vivo* recordings of circadian PER2 expression.

**Fig. 3.**
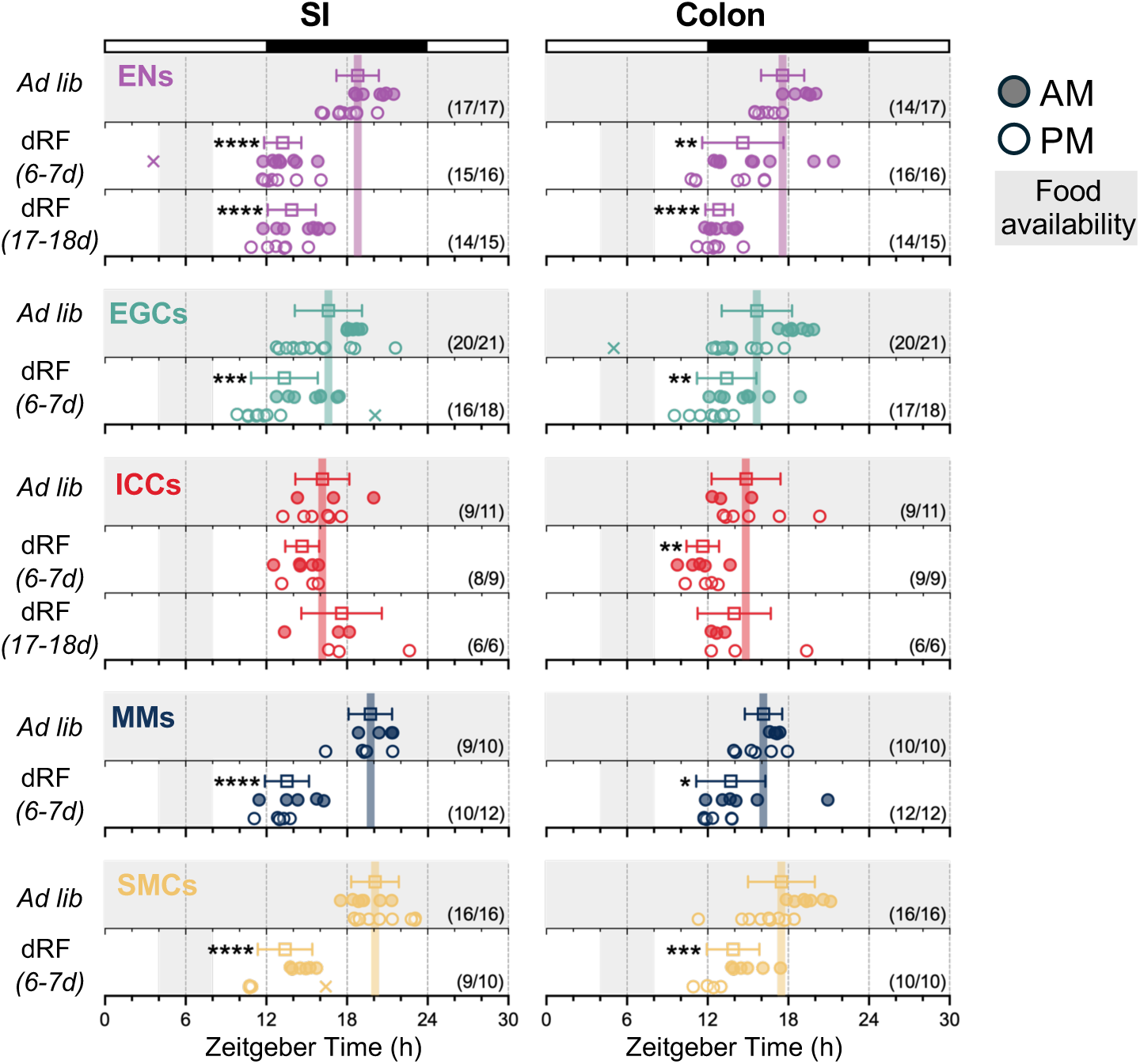
Phase map of five intestinal cell types from mice under *ad libitum* feeding and daytime-restricted feeding. The peak phases of ENs, EGCs, ICCs, MMs, and SMCs from mice fed *ad libitum* (*Ad lib*) or subjected to 6–7 days of daytime-restricted feeding (dRF) are shown. EN and ICC reporters were further examined after long-term dRF (17–18 days). × indicates outliers identified by the ROUT method (Q = 1%); open circles represent AM cultures, and filled circles represent PM cultures. Sample size (shown in the phase plot) reflects the number of tissues with reliably detected rhythms included in the analysis, out of all tissues tested. The gray area indicates food availability. White bar = light; black bar = dark. *p < 0.05, **p < 0.01, ***p < 0.001, ****p < 0.0001; Welch’s t-test.

### Cell-Type–Specific Responses to Food in the Intestinal Clock

It is well-established that feeding can entrain circadian clocks in peripheral organs, including the intestine^35,36^. However, these insights were obtained using the global reporter, which measures aggregate signals from heterogeneous tissues and cannot resolve the rhythms of individual cell types. We therefore used the cell-type–specific PER2 reporting system to examine how *in vivo* manipulation of feeding time impacts the circadian clocks of distinct gut cell types. We compared control mice, fed *ad libitum*, with those on a daytime-restricted feeding (RF) schedule, in which food was only available for 4 hours during the light phase (ZT4–8). This paradigm has been widely used to assess the food-entrainment properties of peripheral clocks^35,37^.

We observed 6–7 days of daytime-RF procedure significantly advanced the phase of PER2::CBG-LUC expression in ENs, SMCs, and MMs by approximately 6 hours (**Fig. 3**), indicating that these cell types can be entrained to a feeding cycle. Interestingly, the magnitude of phase advance was greater in the SI than in the colon, suggesting a segment-dependent difference in food entrainment. To assess the kinetics of food entrainment between segments, EN-specific reporter mice were subjected to daytime-RF for 17–18 days (**Fig. 3**, top plot: ENs). In the SI, there was no further advance following 18 days of daytime-RF, indicating that 6–7 days of daytime-RF is sufficient for SI ENs to fully entrain to the new feeding schedule. In contrast, long-term daytime-RF further advanced the phase of colon ENs, aligning them with those in the SI, suggesting that SI ENs entrain to feeding cues more rapidly than colon ENs.

In contrast, 6–7 days of daytime-RF only modestly affected EGC clocks and had no obvious impact on ICC clocks in the SI (**Fig. 3**), raising the intriguing possibility that the ICC clock may be resistant to altered feeding cycles. To explore this, we subjected ICC-specific reporter mice to daytime-RF for 17–18 days. Strikingly, even after prolonged daytime-RF, the ICC phase was indistinguishable from that observed under *ad libitum* feeding in both the SI and colon (**Fig. 3**, middle plot: ICCs). These results reveal that different intestinal cell types exhibit distinct entrainment properties, a phenomenon that remained elusive prior to the development of our cell-type–specific reporter system and indicate that daytime-RF can induce phase divergence in gut clocks.

### Phase Manipulation of the Enteric Neuronal Clock via In Vivo DREADD Activation

ENs serve as key regulators of complex multicellular communications in the gut^38,39^, but their role in intestinal timekeeping remains unclear. Our data show that circadian clocks in ENs are among the fastest to entrain to feeding schedules, raising the possibility that ENs may function as circadian pacemakers of the intestine. As an initial step to test this idea, we examined whether manipulation of EN activity *in vivo* could alter their intrinsic circadian rhythms. DREADDs are engineered G protein–coupled receptors that can be selectively activated by otherwise inert synthetic ligands, such as Clozapine N-oxide (CNO), enabling precise and reversible control of specific neuronal populations *in vivo*^40^. We aimed to selectively activate ENs *in vivo* using an excitatory DREADD. To achieve this, we crossed R26-LSL-hM3Dq mice, which express the excitatory hM3Dq receptor in a Cre-dependent manner, with *Per2*^CBG-LUC^ mice. Cre-dependent recombination in ENs was induced by intravenous injection of AAV9-CaMKII-Cre, which transduce the majority of ENs (**Supplemental Fig. 2**)^33^, thereby enabling controlled *in vivo* activation of ENs and subsequent *ex vivo* PER2 bioluminescent monitoring of circadian dynamics in DREADD-activated neurons.

For activation of neuronal hM3Dq, we used Compound 21 (C21), a second-generation DREADD ligand designed to avoid CNO’s back-metabolism to clozapine^41^. To test whether daily neuronal activation can shift the circadian phase, C21 was intraperitoneally injected at a time corresponding to the onset of food availability in the daytime-RF paradigm described above. After five days of activation, LMMP from both SI and colon were collected for *ex vivo* circadian monitoring. Although behavioral patterns remained largely unchanged following C21 administration (**Supplemental Fig. 6**), EN activation advanced their phase by approximately 6 hours (**Fig. 4A** and **B**), comparable to the effect of restricted feeding (**Fig. 3**). These observations indicate that the phase shift results directly from neuronal activation rather than from systemic effects. Together, our data demonstrate that EN clocks can be manipulated by direct *in vivo* daily stimulation of ENs. Further study is necessary to determine if these shifts to the phase of EN clocks can cause shifts in the clock phases of other cell types of the gut; to this end, however, our results provide a potential method to investigate this question and dissect the role of ENs in intercellular circadian clock networks within the intestine.

**Fig. 4.**
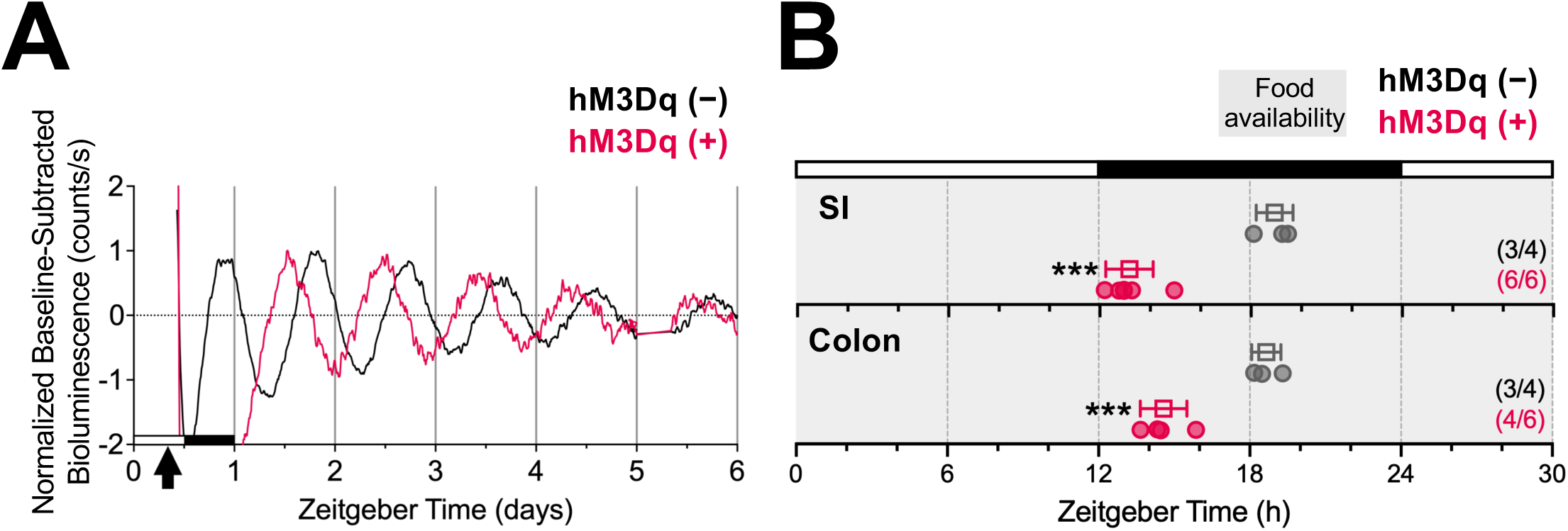
Daytime *in vivo* chemogenetic activation of enteric neurons advances the phase of their circadian rhythm. (A) Representative normalized baseline-subtracted PER2::CBG-LUC traces of the small intestine (SI) of hM3Dq− and hM3Dq+ mice with daily neuronal activation by C21. Baseline-subtracted bioluminescence data were normalized using the first peak after 12 h of recording as 1 (all luminescence values were divided by the value of that peak). Black arrow represents the approximate time when tissue explants were made. (B) Individual peak phases of ENs in SI and colon LMMP explants from hM3Dq− and hM3Dq+ mice with daily intraperitoneal C21 injections. Sample size (shown in the phase plot) reflects the number of tissues with reliably detected rhythms included in the analysis, out of all tissues tested. ***p < 0.001; Welch’s t-test.

## Discussion

Circadian clocks are ubiquitous, and nearly all cells in central and peripheral adult organs contain cell-autonomous oscillators. Recent single-cell and single-nucleus RNA sequencing studies have demonstrated that distinct cell types within the same tissues can exhibit unique circadian phases^42,43^. In the liver and white adipose tissue, most rhythmically-expressed genes are entrained by feeding cycles. In contrast, in brown adipose tissue and skin, only a subset of rhythmic transcripts shifts with feeding while others retain their original phase^44,45^. These observations indicate that cellular circadian organization is highly heterogeneous and that individual cell types may differentially respond to extracellular cues such as feeding. Therefore, cell-type–specific circadian reporters are required to understand the organization and entrainment of individual cellular clocks within heterogeneous tissues.

To the best of our knowledge, only two Cre-dependent circadian reporter mouse lines have previously been generated^46,47^. Previously, we (Y.S. and J.S.T.) generated Color-Switch PER2::LUC mice, in which PER2 is translated as a fusion protein with click beetle green luciferase (CBG-LUC) in Cre-positive cells and with click beetle red luciferase (CBR-LUC) in Cre-negative cells^46^. Using an ultra-sensitive, low-light detection camera, it is possible to simultaneously record single-cell bioluminescence rhythms from both Cre-positive and Cre-negative cells in tissues explanted from Color-Switch PER2::LUC mice. However, no commercially available bioluminescence imaging system currently provides the sensitivity required for dual color single-cell recording. Building such a system is costly and requires extensive expertise in optics and imaging, and even then, it can typically record only one sample at a time. Although single-cell imaging has distinct advantages, in most applications it is sufficient to record population-level rhythms from specific cell types within a tissue. Another Cre-dependent circadian reporter is the *Dbp*-LUC knock-in mouse^47^, which enables both *in vivo* and *ex vivo* detection of circadian bioluminescence rhythms in numerous tissues. However, because this reporter uses firefly luciferase, its signal is weaker than that of CBG-LUC and may fall below the detection threshold when only a small number of cells within an explant express the reporter. In contrast, the *Per2^CBG-LUC^* reporter mice developed in the present study are well suited for measuring circadian bioluminescence from relatively small populations of cells within complex multicellular organs using standard, commercially-available bioluminescence monitoring systems without any modifications.

Intestinal tissues comprise diverse cell types capable of responding to environmental changes and communicating extensively with each other to coordinate physiological processes. In particular, gut-intrinsic ENs, regarded as central coordinators of intestinal tissue circuits^39^, are strategically positioned to sense environmental cues, such as dietary components, and translate them into coordinated physiological responses, including motility, secretion, and immunity^38,48^. Our data indicate that circadian clocks in ENs are among the fastest to entrain to feeding schedules. Given their role in communicating with other intestinal cell types, these findings raise the possibility that ENs may “phase-lead” in coordinating circadian rhythms across gut cell populations. Supporting this notion, the clocks of SMCs, a cell type directly regulated by ENs to modulate intestinal peristalsis, exhibit circadian phases similar to those of ENs under both steady-state conditions and in response to daytime-restricted feeding. Although signaling from ENs to MMs is only beginning to be explored^30,49,50^, the similar behavior of their clocks suggests that ENs may also influence MM clocks, contributing to the regulation of tissue homeostasis and immune function at the appropriate times of day. In contrast, EGCs and ICCs may use entrainment mechanisms distinct from those of ENs, as reflected by their differing circadian phase patterns. Future studies employing neuron-specific manipulation of circadian clocks will help determine their influence on the circadian phases of SMCs and MMs. To support the feasibility of this approach, our experiments demonstrated that *in vivo* DREADD activation of neurons, including ENs, significantly advanced the circadian phase of ENs without altering behavior. These experimental strategies will help determine whether a hierarchical system exists in which a specific cell type, such as ENs, functions as a circadian pacemaker for the entire gut, introducing a novel framework for understanding circadian organization within multicellular organs.

Circadian regulation is a fundamental determinant of intestinal health^14–19^. Many individuals experience intestinal disturbances, such as constipation, when undergoing circadian disruptions, for example, due to travel across time zones^51,52^. Beyond these temporary discomforts, growing evidence indicates that circadian disruption can profoundly influence the development of, and susceptibility to, major intestinal disorders, including inflammatory bowel disease (IBD) and disorders of gut-brain interactions (DGBIs), such as irritable bowel syndrome (IBS)^14–19^. Moreover, recent studies in mice have partially recapitulated these pathological changes^53–55^. Mice subjected to a shifted light-dark cycle exhibit increased intestinal barrier permeability^55^, altered microbiota composition^53^, and enhanced susceptibility to chemically-induced colitis^54^. However, it remains unclear how circadian misalignment affects the cellular network of gut clocks. Our data indicate that intestinal cell types respond differently to altered feeding schedules: ENs, SMCs, and MMs exhibit rapid phase shifts, whereas EGCs and ICCs are comparatively resistant. These findings establish time-restricted feeding as a useful paradigm for exploring how divergence in gut clocks may contribute to intestinal homeostasis and disease susceptibility. The observation of heterogeneous phase shifts also suggests that intestinal clocks in distinct cell types may re-entrain at different rates following light-dark shifts. Future studies will quantify re-entrainment kinetics in response to altered light-dark cycles and examine how jet-lag or shift-work paradigms disrupt circadian organization among different gut cell types. By linking cell-specific circadian dynamics to functional outcomes, this approach provides new insights into how intestinal clocks respond to environmental changes to maintain gut homeostasis.

Our findings also reveal distinct rates of food entrainment in ENs between the SI and colon. ENs in the SI, where nutrient absorption occurs, appear highly sensitive to feeding cues, potentially facilitating circadian coordination of digestive processes. In contrast, ENs in the colon exhibit slower food entrainment, which may reflect colon-specific adaptations in which microbial regulation plays a more dominant role than dietary cues. Indeed, recent studies have reported region-specific transcriptional differences in ENs between SI and colon, shaped in part by local environmental factors, including the microbiota^33,56^. Future studies will investigate whether altered composition of microbiota affects the efficiency of colonic EN entrainment to feeding schedules. Our findings on region-specific differences in food entrainment may inform the design of circadian interventions, such as the timing of meals or probiotic administration, to more effectively target intestinal function and promote gut health.

Taken together, our data highlight: (1) cell-type- and region-specific dynamics of circadian gene expression in gut clocks; (2) their distinct responsiveness to feeding cues; and (3) a technical advance that enables simultaneous manipulation and monitoring of the circadian clock in ENs. This approach provides a new avenue for elucidating the molecular basis of gut clock entrainment and, when combined with *in vivo* manipulations (e.g., daytime-restricted feeding, or shifted light-dark cycle), will allow investigation of which signals, such as light-induced brain-body cues or food intake, individual intestinal cells utilize as their primary entraining factors. Defining gut circadian clock networks is essential for understanding intestinal health and may pave the way for developing strategies to prevent functional impairments in intestinal physiology caused by circadian disruption.

## Materials and Methods

### Animals

All mice were maintained under a 12:12 light dark (LD) cycle in SPF barrier or conventional facilities at the University of Texas Southwestern Medical Center. The PER2::CBG-LUC knock-in mouse line (*Per2*^CBG-LUC^) was created as described in **Supplemental Fig. 1** and maintained in a C57BL/6N background. *Per2*^CBG-LUC^ mice were crossed with E2a-Cre (JAX #:003724) or with one of five cell-type–specific Cre drivers to target enteric neurons (BAF53b-Cre, JAX#027826), muscularis macrophages (Csf1r-Cre, JAX#029206), enteric glial cells (PLP1b-CreERT2, JAX#005975), interstitial cells of Cajal (cKit-CreERT2), or smooth muscle cells (Myh11-CreERT2-RAD, JAX#037658), all on a C57BL/6NJ mixed background. For DREADD-based chemogenetic activation of neurons *in vivo*, *Per2*^CBG-LUC^ mice were crossed with R26-LSL-hM3Dq-DREADD (JAX# 026220) on C57BL/6NJ mixed background. cKit-CreERT2 was kindly provided by Dr. Dieter Saur (Technical University of Munich)^29^ through the lab of Drs. Kenton M. Sanders and Sal Baker (University of Nevada).

Induction of CreERT2 activity was achieved by intraperitoneal injection of tamoxifen (20 mg/mL; Sigma-Aldrich, T5648) dissolved in corn oil (Sigma-Aldrich, C8267). Mice received injections on two consecutive days at a dose of 0.1 mg per g body weight, 14 days prior to isolation of intestinal tissues. Cre targeting of enteric neurons was also achieved by intravenous injection of *Per2*^CBG-LUC^ mice with AAV9-CaMKII-Cre (Addgene #105558) as previously described^33^. All mice used in this study were heterozygous for both the PER2::CBG-LUC knock-in allele and the Cre transgene and were between 6 and 23 weeks of age at the time of experiments.

Genotyping was performed by Transnetyx, and probe information is available from Transnetyx or upon request. We occasionally observed germline Cre recombination BAF53b-Cre line crossed with *Per2*^CBG-LUC^ mice. To identify germline-recombined animals, we collected liver and white adipose tissues at the time of gut explant preparation and performed acute luminescence measurements. If ectopic luminescent signals were detected in these tissues, the corresponding gut explant data were excluded from analysis. Germline Cre recombination could also be judged from abnormally high luminescent intensity observed in gut explants.

Mice maintained under *ad libitum* feeding conditions (*Ad lib*) were housed in the barrier animal facility. For daytime-restricted feeding and DREADD experiments, mice were individually housed in cages (32.5 cm in length, 14.5 cm in width, and 13 cm in height, equipped with an 11-cm diameter running wheel) with woodchip bedding (Sani-Chips, PJ Murphy Forest Products, Montville, NJ, USA). The cages were placed in light-tight ventilated cabinets. Mice were under a 12 h light (∼200 lux at cage level, generated by white LEDs at the top of the light-tight box) and 12 h dark condition, and the light condition was controlled by the Chamber Controller software (ver. 4.104, Actimetrics, Lafayette, IN, USA). Wheel-running activity (collected by micro-switches mounted to the sides of the running wheel cages) was continuously recorded every 1 minute by the ClockLab acquisition system (version 3.604, Actimetrics, Lafayette, IN, USA) for each mouse. Cages and water bottles were changed every 3 weeks. After several days of baseline behavioral recording under *ad lib* feeding to monitor circadian activity rhythms, animals were subjected to an overnight fasting period. Daytime-restricted feeding was then implemented sequentially according to the following schedule: 8-hour feeding for 2 days, 6-hour feeding for 2 days, and 4-hour feeding for either short-term (6–7 days) or long-term (17–18 days) experiments. Food was provided daily at Zeitgeber Time (ZT) 4–8 manually. For DREADD-based chemogenetic activation of neurons *in vivo*, mice were individually housed in running-wheel cages with *ad lib* access to chow and subjected to the chemogenetic activation protocol described below (see *In vivo* DREADD section). All animal procedures were approved by the Institutional Animal Care and Use Committee (IACUC) of the University of Texas Southwestern Medical Center (APN 2016-101376 and APN 2021-103099).

### Generation of PER2::CBG-LUC knock-in mice

The PER2::CBG-LUC knock-in targeting vector was derived from the backbone of the PER2::LUC targeting vector^6^. Firefly luciferase was replaced with a reporter cassette consisting of a floxed NeoR and green click beetle luciferase (CBG)^57^. The CBG open reading frame (ORF) was fused to the read-through frame of the second loxP site (11 aa) and positioned in conjunction with the 3’ end of the *Per2* ORF in exon 23. To minimize non-specific expression of CBG prior to Cre-Lox recombination, the ATG start codon was removed and three stop codons were inserted immediately upstream of the CBG ORF. The 24-amino-acid (72 bp) truncation present in the previous version of the PER2::LUC reporter was corrected to restore a full-length PER2 ORF^6^.

ES cell culture and gene targeting were performed as described previously^46^. Genomic DNA from screened ES cells was prepared, and long-fragment PCR (LA Taq, Takara) and Southern blotting^6^ were carried out to verify integration of the non-genomic DNA sequence at the *Per2* locus. Targeted ES cells were injected into B6(Cg)-Tyrc−2J/J blastocysts and transferred to pseudopregnant B6(Cg)-Tyrc−2J/J female recipients. Resulting male chimeras were bred with C57BL/6N females, and germline transmission was confirmed by coat color, bioluminescence, and Southern blotting.

### Isolation and immunofluorescence staining of longitudinal muscle–myenteric plexus (LMMP)

Wholemount staining of LMMP tissues was performed as previously described^58^. Briefly, intestines were prepared by first removing adipose tissues and flushing the luminal contents with cold PBS. A 1-mL pipette tip was inserted into the intestinal lumen to fully extend the intestinal wall. The LMMP layer was then separated from the mucosa using cotton swabs, as previously described^24^. The LMMP was stretched and pinned onto a Sylgard-coated Petri dish (Fisher Scientific) containing cold PBS, and then fixed in 4% PFA overnight at 4 °C.

Following fixation, tissues were rinsed five times with PBS at room temperature with gentle shaking, then permeabilized and blocked in PBS containing 1% Triton X-100 and 10% normal donkey serum for 1 hour at room temperature. Samples were incubated with primary antibodies in the same buffer overnight at 4 °C, washed with PBS containing 1% Triton X-100, and then incubated with secondary antibodies in blocking buffer for 3 hours at room temperature. Immunostained tissues were washed four times with PBS containing 1% Triton X-100, and after a final PBS rinse, mounted on Superfrost microscope slides using VECTASHIELD (Vector Laboratories).

Fluorescently labeled LMMP preparations were imaged using a spinning disk confocal microscope (Nikon) equipped with a Hamamatsu Orca-Fusion sCMOS camera and controlled by NIS-Elements Advanced Research software (Nikon). All image analyses were performed using the image-processing package Fiji.

### Tissue explants and bioluminescence recording

Mice were euthanized by cervical dislocation followed by decapitation without anesthesia at two distinct time points, ZT2–4 and ZT10–13 (ZT0 = lights-on). The ileum and proximal colon were dissected after removing surrounding adipose tissue. A tip of 1-mL pipette was inserted into the lumen to extend the intestinal wall, and the longitudinal muscle–myenteric plexus (LMMP) layer was separated from the mucosa using cotton swabs^24^.

LMMP sheets (approximately 2 cm in length; wet weight, 7–10 mg for small intestine and 10–15 mg for colon) were rinsed three times in cold, HEPES-buffered HBSS containing antibiotics (500 mL HBSS H9394-500ML, 5 mL of 1M HEPES solution H0887, 5 mL of Penicillin-Streptomycin solution P0781; Sigma-Aldrich). Ileal and colonic LMMP explants were cultured on 500-µm woven filters (Spectra/Mesh 146418; Repligen) in 1.5 mL of B27-supplemented recording medium containing luciferin. 1 L recording media contained 12.77g of DMEM (90-013PB, CellGro), 20 mL of L-glutamine solution (G7513, Sigma-Aldrich), 4.7 mL of 7.5% sodium bicarbonate solution (S8761, Sigma-Aldrich), 10 mL of 1M HEPES solution (H0887, Sigma-Aldrich), and 2.5 mL of Penicillin-Streptomycin solution (P0781, Sigma-Aldrich). Osmolality of recording media typically range between 273 and 275 mOsm. B27 supplement (1 mL for 50 mL, 17504-044, Gibco) and beetle luciferin potassium salt (0.1mM final concentration, E160B, Promega) were added a few days before explants were made. Each culture was sealed by vacuum grease and covered with a 40-mm diameter circle cover glass. Manual isolation of the LMMP layers in both the SI and colon was completed within a few minutes for each sample with a total preparation time of approximately 10 minutes per mouse (from euthanasia to the start of luminescence recording for both tissues). This rapid processing was essential to prevent tissue damage and circadian phase resetting by culture procedure. Bioluminescence was recorded continuously using the LumiCycle 32 (Actimetrics) with specially hand-selected low dark-count photomultiplier photon-counting heads (H6240MOD, Hamamatsu). Total photon number emitted from each tissue was counted for 1 min every 10 min interval. Bioluminescence was recorded for 3–7 days.

Bioluminescent data were analyzed using LumiCycle Analysis software (v3.101; Actimetrics). Raw traces were baseline-corrected using a 24-h moving average and smoothed with a 0.5-h adjacent average. Since early peaks, including the first peaks for AM cultures, were often masked by rapid changes of the baseline, the circadian phase was defined as the second peak for AM cultures and the first peak detected after 12 h in culture for PM cultures. Only samples in which the circadian phase could be reliably detected were included in the analysis, and the number of analyzed samples relative to the total collected is indicated in the figure.

### In vivo DREADD activation

For chemogenetic activation and PER2::CBG-LUC monitoring of neurons, Rosa26-LSL-hM3Dq-DREADD mice were crossed with *Per2*^CBG-LUC^ reporter mice. The resulting mice were intravenously injected with AAV9-CaMKII-Cre three weeks prior to tissue collection. C21, a selective DREADD agonist, was administered intraperitoneally at a dose of 3 mg/kg body weight^59^, once daily at Zeitgeber Time (ZT) 4 for five consecutive days. On the fifth day of C21 administration, mice were euthanized, and small intestinal and colonic LMMP tissues were collected for ex vivo bioluminescence recording.

### Statistical analysis

All statistical analyses were performed using GraphPad Prism (version 10.6.1; Dotmatics, Boston, MA, USA; RRID:SCR_002798). Outliers were identified using the ROUT test (Q = 1%). Differences in phase were assessed using Welch’s t-test. Statistical significance was defined as p < 0.05 (*p < 0.05, **p < 0.01, ***p < 0.001, ****p < 0.0001). Detailed statistical methods and exact p-values are provided in the figure legends.

## Key Resources

| REAGENT or RESOURCE | SOURCE | IDENTIFIER |
| --- | --- | --- |
| **Antibodies** |  |  |
| Rabbit monoclonal anti-HuC/HuD | Abcam | Cat# ab184267 |
| Rabbit polyclonal anti-S100 | Dako Omnis | Cat# GA504 |
| Rabbit polyclonal anti-ACTA2 | ABclonal | Cat# A1011 |
| Goat polyclonal anti-CD117/c-Kit | R&D Systems | Cat# AF1356 |
| Rat monoclonal anti-F4/80 | Invitrogen | Cat# 14-4801-82 |
| Alexa Fluor® 594 AffiniPure® Donkey Anti-Rabbit IgG (H+L) | Jackson ImmunoResearch | Cat# 711-585-152 |
| Alexa Fluor® 647 AffiniPure® Donkey Anti-Goat IgG (H+L) | Jackson ImmunoResearch | Cat# 705-605-147 |
| Rhodamine (TRITC) AffiniPure® Donkey Anti-Rat IgG (H+L) | Jackson ImmunoResearch | Cat# 712-025-150 |
| **Bacterial and virus strains** |  |  |
| AAV9-CamKII-Cre (6.11x10<sup>13</sup> GC/mL) | VectorBuilder | VB231127-1512psf |
| **Chemicals, peptides, and recombinant proteins** |  |  |
| DMEM | Corning-Cellgro | 90-013-PB |
| Beetle Luciferin Potassium Salt | Promega | E160B |
| B-27 Supplement | Gibco | 17504-044 |
| L-Glutamine solution | Sigma-Aldrich | G7513 |
| HBSS | Sigma-Aldrich | H9394 |
| HEPES | Sigma-Aldrich | H0887 |
| Sodium Bicarbonate | Sigma-Aldrich | S8761 |
| Penicillin-Streptomycin | Sigma-Aldrich | P0781 |
| Normal Donkey Serum | SouthernBiotech | Cat# 0030-01 |
| Triton X-100 | Alfa Aesar | Cat# A16046.AP |
| Dulbecco's Phosphate-Buffered Saline, 1X without calcium and magnesium | Corning | Cat# 21-031-CV |
| DREADD agonist 21 dihydrochloride | Tocris Bioscience | Cat# 6422 |
| Tamoxifen | Sigma-Aldrich | Cat# T5648 |
| Corn oil | Sigma-Aldrich | Cat# C8267 |
| **Experimental models: Organisms/strains** |  |  |
| Mouse: Per2::CBG-LUC | This paper |  |
| Mouse: STOCK Tg(Actl6b-Cre)4092Jiwu/J | The Jackson Laboratory | JAX: 027826 |
| Mouse: B6.Cg-Tg(Plp1-cre/ERT)3Pop/J | The Jackson Laboratory | JAX: 005975 |
| Mouse: C57BL/6-Tg(Myh11-cre/ERT2)F31Gko/J | The Jackson Laboratory | JAX: 037658 |
| Mouse: C57BL/6-Tg(Csf1r-cre)1Mnz/J | The Jackson Laboratory | JAX: 029206 |
| Mouse: 129.SvTac-Kittm1(cre/ERT2)Dsa | Klein et al. | MGI: 5543260 |
| Mouse: B6.FVB-Tg(Ella-cre)C5379Lmgd/J | The Jackson Laboratory | JAX: 003724 |
| Mouse: B6.129-Gt(ROSA)26Sortm5.1(CAG-Sun1/sfGFP)Nat/MmbeJ | The Jackson Laboratory | JAX: 030952 |
| Mouse: B6N;129-Tg(CAG-CHRM3*, -mCitrine)1Ute/J | The Jackson Laboratory | JAX: 026220 |
| **Software and algorithms** |  |  |
| ImageJ (v. 2.14.0/1.54f) | National Institutes of Health | https://imagej.nih.gov/ij/ |
| LumiCycle Acquisition Software (version 5.01) | Actimetrics, Inc. | https://actimetrics.com/products/lumicycle/ |
| LumiCycle Analysis Software (version 3.101) | Actimetrics | https://actimetrics.com/products/clocklab/ |
| ClockLab Acquisition Software (version 3.604) | Actimetrics | https://actimetrics.com/products/clocklab/ |
| Chamber Controller Software (version 4.104) | Actimetrics | https://actimetrics.com/products/clocklab/ |
| ClockLab Analysis Software (version 6.1.18) | Actimetrics | https://actimetrics.com/products/clocklab/ |
| Prism | GraphPad Software | Version 10.5.0 |
| **Other** |  |  |
| Stereo dissection microscope | Leica | MZ6 |
| 0.5 mm Woven Filter | Spectra/Mesh | 146418 |
| Cover Glass 40 mm Circle #1 | Fisher | 12541011 |
| High vacuum grease | Dow Corning | NA |
| 6-inch Cotton Swab | Amazon | NA |
| 28G Lo-Dose™ U-100 Insulin Syringes | BD | 329461 |
| Rodent Chow | Harlan | Teklad Global 18% Protein Rodent Diet 2918 |

## ACKNOWLEDGMENTS

We thank Aliza Abbas, Sofia Farah, Emily John, and Avalene Tan for their valuable discussions and experimental support. We also acknowledge the Transgenic Core at UT Southwestern directed by Dr. Robert E. Hammer and the Quantitative Light Microscopy Core (QLMC) directed by Dr. Marcel Mettlen for their technical assistance.

This work was supported by grants from the National Institutes of Health R01NS114527 awarded to S.Y. , R01NS106657 awarded to J.S.T., and the National Science Foundation IOS-1931115 awarded to S.Y. Y. O. is supported by the Nancy Cain and Jeffrey A. Marcus Scholar Award in Medical Research (in honor of B. S. Vowell), the Pew Scholars Program in Biomedical Sciences (00036890), and the Welch Foundation Research Grant (I-2153-20230405). J.S.T. was an Investigator in the Howard Hughes Medical Institute.

## AUTHOR CONTRIBUTIONS

Y.O. and S.Y. conceived and designed the research. Y.S. and J.T. generated the *Per2*^CBG-LUC^ mouse line. Y.O., I.M., and S.Y. performed experiments with support from S.K.T.T. and D.E. I.M., M.S., S.K.T.T., Y.O., and S.Y. analyzed the data and interpreted the results. Y.O. drafted the manuscript with inputs from S.Y., S.K.T.T., and M.S. M.S. and Y.S. generated some schematic illustrations. All authors participated in editing the manuscript and have approved the final version.

The authors declare no competing interests

---

**Supplemental Fig. 1.**
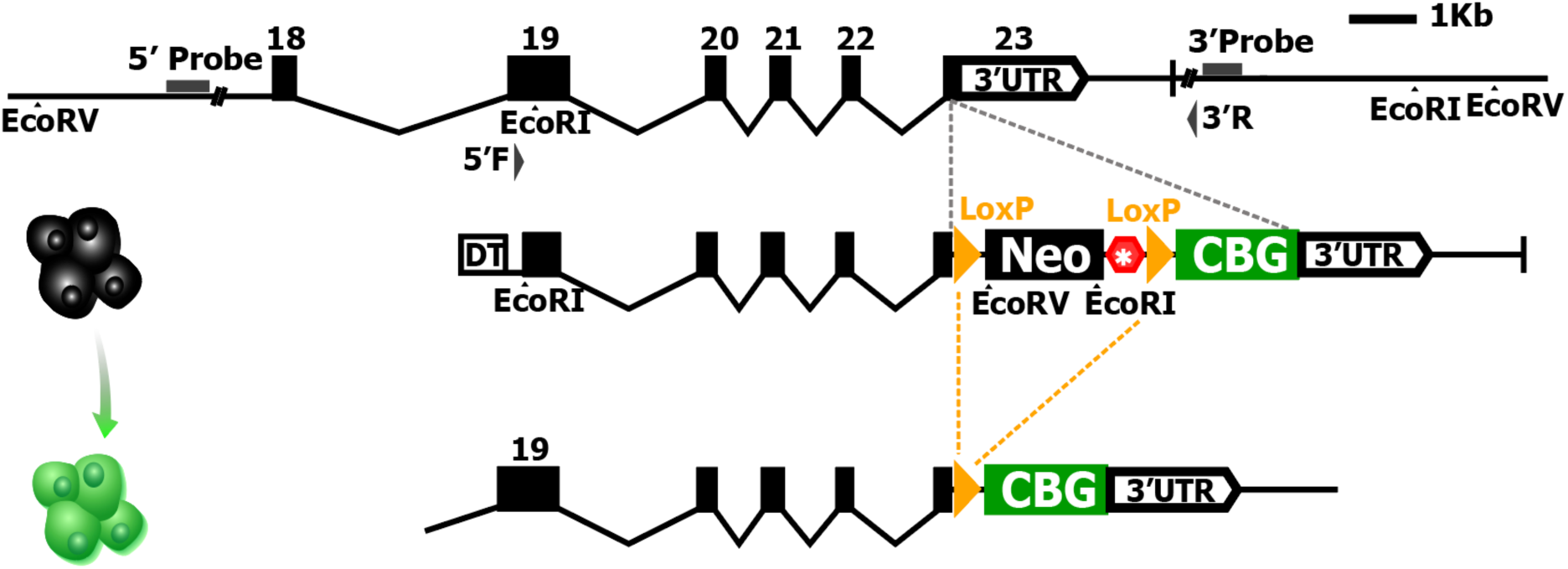
Diagram of the PER2::CBG-LUC knock-in construct targeted to the *Per2* locus. Yellow triangles indicate the LoxP sites flanking the STOP cassette. Cre-mediated recombination excises the STOP cassette, allowing PER2::CBG-LUC expression specifically in Cre-expressing cells.

**Supplemental Fig. 2.**
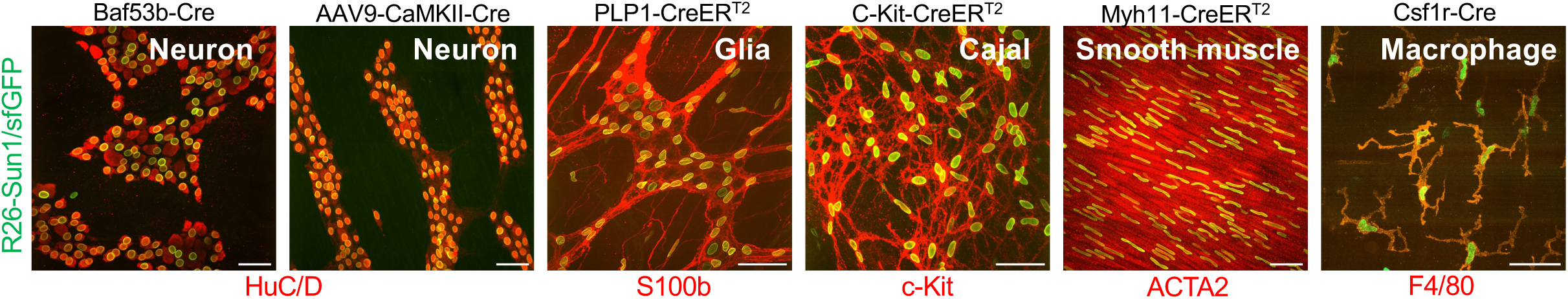
Cre-dependent targeting of various intestinal cell types in the muscularis externa. Representative colon LMMP tissues stained with cell-type–specific markers are shown: HuC/D (ENs), S100b (EGCs), c-Kit (ICCs), ACTA2 (SMCs), and F4/80 (MMs). Red: marker staining; Green: nuclear-localized GFP reporter activated by Cre recombination. Scale bars: 50 μm. The R26-Sun1/sfGFP reporter was crossed with each Cre driver or intravenously injected with AAV9-CaMKII-Cre. Images were taken from at least five distinct regions of LMMP tissues. Data were collected from 2 mice per reporter line.

**Supplemental Fig. 3.**
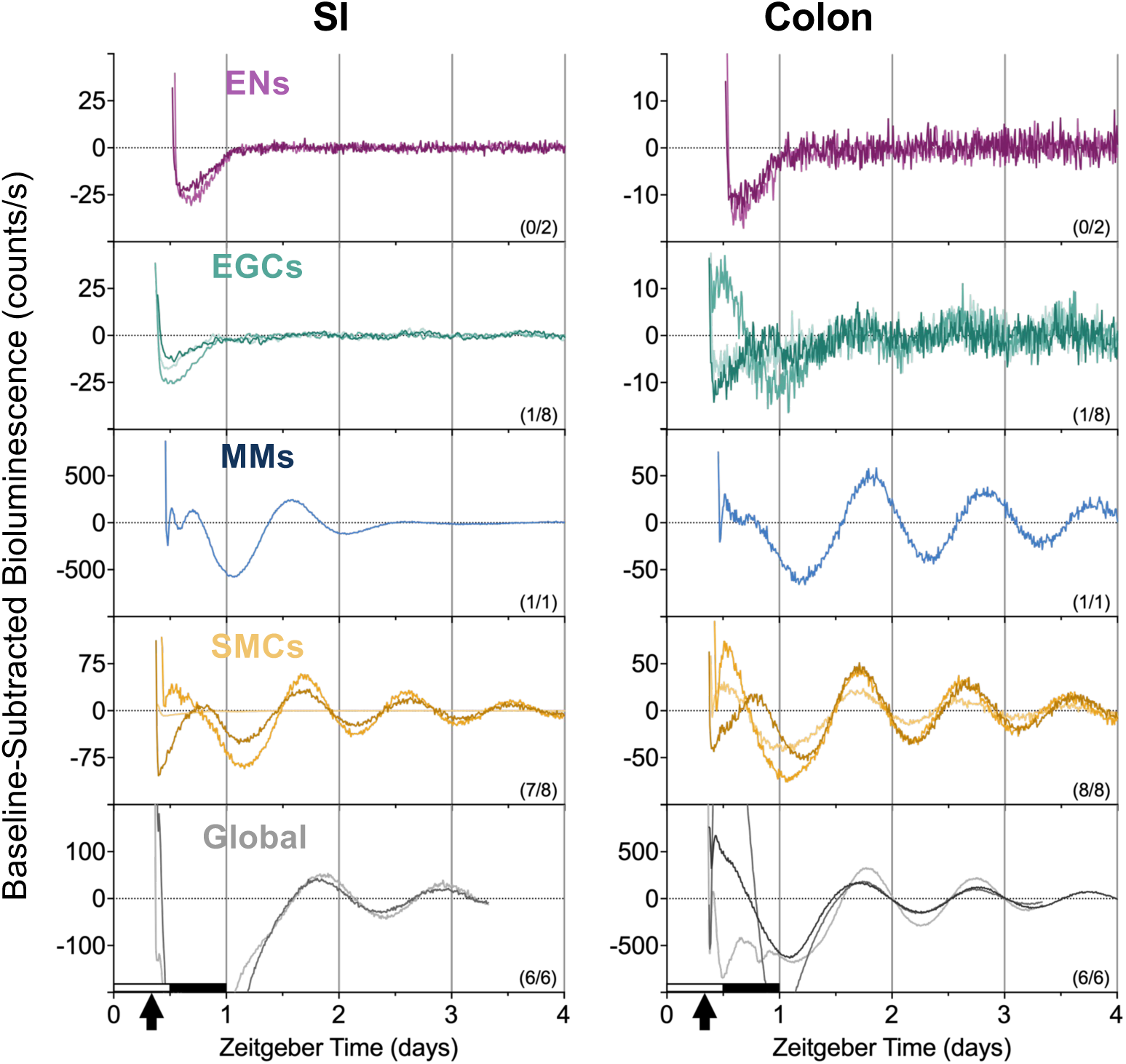
Representative baseline-subtracted PER2::CBG-LUC traces of ENs, EGCs, MMs, SMCs, and all cells in submucosal layer. Sample size (shown in the plot) reflects the number of tissues with reliably detected rhythms included in the analysis, out of all tissues tested. Black arrows represent the approximate time when tissue explants were made.

**Supplemental Fig. 4.**
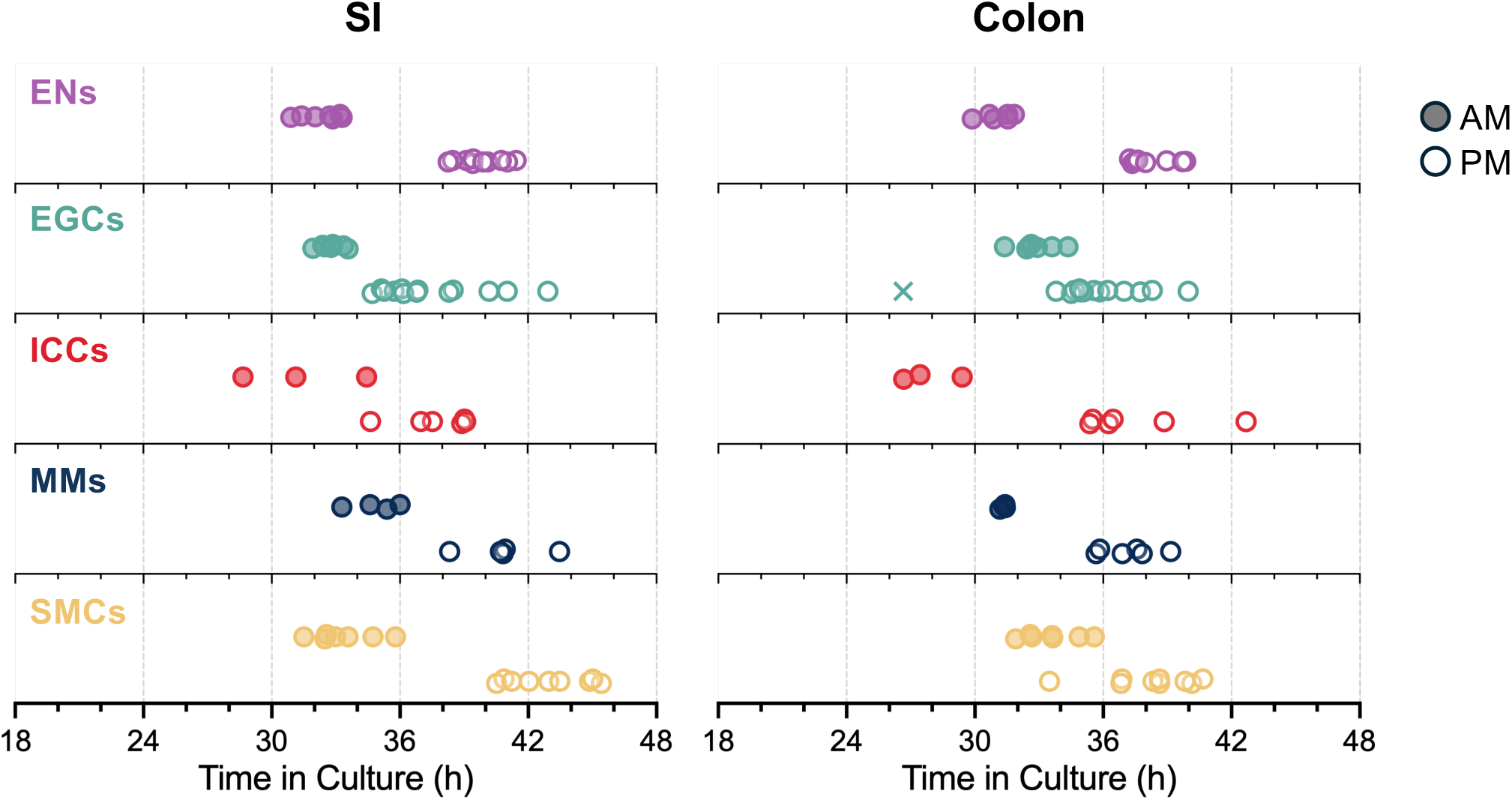
Peak phases of individual tissue explants plotted relative to culture time. The same data plotted in Fig. 3 (plotted relative to Zeitgeber Time) were plotted relative to culture time. × indicates outliers identified by the ROUT method (Q = 1%); open circles represent AM cultures, and filled circles represent PM cultures.

**Supplemental Fig. 5.**
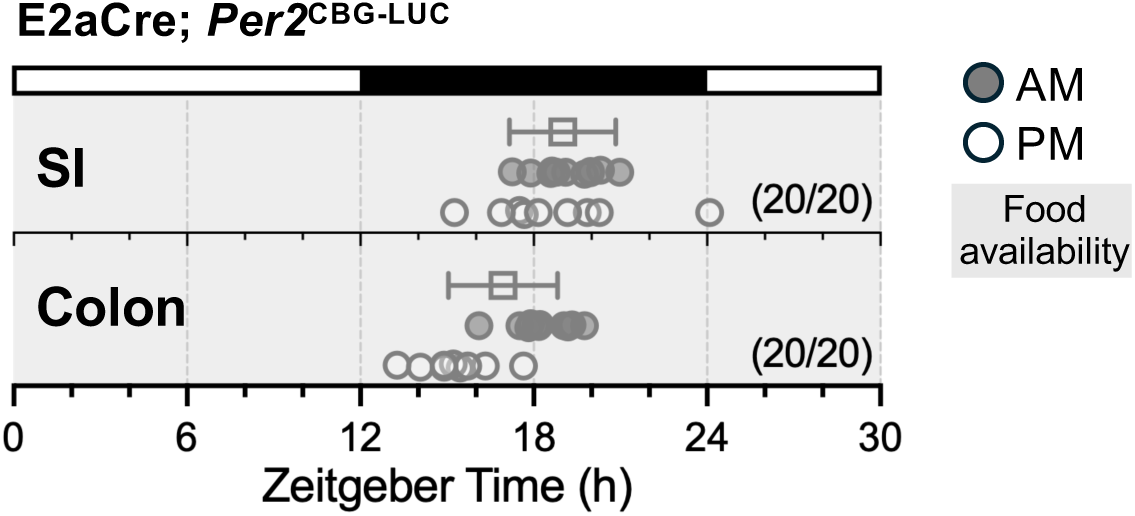
Phase map of gut explants from *E2a-Cre; Per2^CBG-LUC^* mice. The peak phase of small intestine (SI) and colon from tissue explanted from mice under *ad libitum* feeding. Open circles represent AM cultures, and filled circles represent PM cultures. Sample size (shown in the phase plot) reflects the number of tissues with reliably detected rhythms included in the analysis, out of all tissues tested. The gray area indicates food availability. White bar = light; black bar = dark.

**Supplemental Fig. 6.**
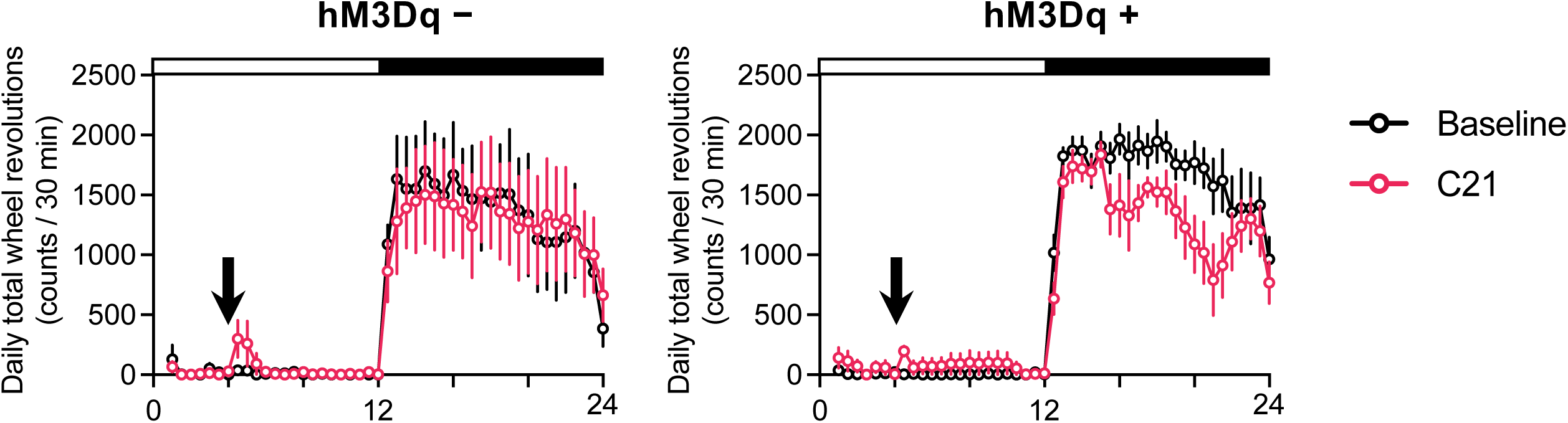
Daily C21 injection had negligible impact on behavioral activity rhythm. Group-averaged 24-h activity profiles (mean ± SEM) during baseline (4 days before C21 injection; black) and during C21 injection (red) are shown. Tissues were explanted on the evening of the fifth C21 injection day. C21 treatment did not induce anticipatory behavior, although a modest decrease in mid-night wheel-running activity was observed (right). n = 6. C21, DREADD agonist.

## REFERENCES

1. Vitaterna, M.H., Takahashi, J.S., and Turek, F.W. (2001). Overview of circadian rhythms. Alcohol Res Health 25, 85–93.

2. Takahashi, J.S. (2004). Finding new clock components: past and future. J Biol Rhythms 19, 339–347. 10.1177/0748730404269151.

3. Whitmore, D., Sassone-Corsi, P., and Foulkes, N.S. (1998). PASting together the mammalian clock. Curr Opin Neurobiol 8, 635–641. 10.1016/s0959-4388(98)80092-5.

4. Takahashi, J.S. (2017). Transcriptional architecture of the mammalian circadian clock. Nat Rev Genet 18, 164–179. 10.1038/nrg.2016.150.

5. Yamazaki, S., Numano, R., Abe, M., Hida, A., Takahashi, R., Ueda, M., Block, G.D., Sakaki, Y., Menaker, M., and Tei, H. (2000). Resetting central and peripheral circadian oscillators in transgenic rats. Science 288, 682–685. 10.1126/science.288.5466.682.

6. Yoo, S.H., Yamazaki, S., Lowrey, P.L., Shimomura, K., Ko, C.H., Buhr, E.D., Siepka, S.M., Hong, H.K., Oh, W.J., Yoo, O.J., et al. (2004). PERIOD2::LUCIFERASE real-time reporting of circadian dynamics reveals persistent circadian oscillations in mouse peripheral tissues. Proc Natl Acad Sci U S A 101, 5339–5346. 10.1073/pnas.0308709101.

7. Davidson, A.J., Yamazaki, S., and Menaker, M. (2003). SCN: ringmaster of the circadian circus or conductor of the circadian orchestra? Novartis Found Symp 253, 110–121; discussion 121-115, 281-114.

8. Brown, A.J., Pendergast, J.S., and Yamazaki, S. (2019). Peripheral Circadian Oscillators. Yale J Biol Med 92, 327–335.

9. Mohawk, J.A., Green, C.B., and Takahashi, J.S. (2012). Central and peripheral circadian clocks in mammals. Annu Rev Neurosci 35, 445–462. 10.1146/annurev-neuro-060909-153128.

10. Bishehsari, F., Post, Z., Swanson, G.R., and Keshavarzian, A. (2025). Circadian Rhythms in Gastroenterology: The Biological Clock’s Impact on Gut Health. Gastroenterology. 10.1053/j.gastro.2025.06.017.

11. Zheng, D., Ratiner, K., and Elinav, E. (2020). Circadian Influences of Diet on the Microbiome and Immunity. Trends Immunol 41, 512–530. 10.1016/j.it.2020.04.005.

12. Segers, A., and Depoortere, I. (2021). Circadian clocks in the digestive system. Nat Rev Gastroenterol Hepatol 18, 239–251. 10.1038/s41575-020-00401-5.

13. Brooks, J.F., 2nd, Behrendt, C.L., Ruhn, K.A., Lee, S., Raj, P., Takahashi, J.S., and Hooper, L.V. (2021). The microbiota coordinates diurnal rhythms in innate immunity with the circadian clock. Cell 184, 4154–4167 e4112. 10.1016/j.cell.2021.07.001.

14. Nojkov, B., Rubenstein, J.H., Chey, W.D., and Hoogerwerf, W.A. (2010). The impact of rotating shift work on the prevalence of irritable bowel syndrome in nurses. Am J Gastroenterol 105, 842–847. 10.1038/ajg.2010.48.

15. Kim, H.I., Jung, S.A., Choi, J.Y., Kim, S.E., Jung, H.K., Shim, K.N., and Yoo, K. (2013). Impact of shiftwork on irritable bowel syndrome and functional dyspepsia. J Korean Med Sci 28, 431–437. 10.3346/jkms.2013.28.3.431.

16. Ananthakrishnan, A.N., Khalili, H., Konijeti, G.G., Higuchi, L.M., de Silva, P., Fuchs, C.S., Richter, J.M., Schernhammer, E.S., and Chan, A.T. (2014). Sleep duration affects risk for ulcerative colitis: a prospective cohort study. Clin Gastroenterol Hepatol 12, 1879–1886. 10.1016/j.cgh.2014.04.021.

17. Zhen Lu, W., Ann Gwee, K., and Yu Ho, K. (2006). Functional bowel disorders in rotating shift nurses may be related to sleep disturbances. Eur J Gastroenterol Hepatol 18, 623–627. 10.1097/00042737-200606000-00008.

18. Ananthakrishnan, A.N., Long, M.D., Martin, C.F., Sandler, R.S., and Kappelman, M.D. (2013). Sleep disturbance and risk of active disease in patients with Crohn’s disease and ulcerative colitis. Clin Gastroenterol Hepatol 11, 965–971. 10.1016/j.cgh.2013.01.021.

19. Knutsson, A., and Boggild, H. (2010). Gastrointestinal disorders among shift workers. Scand J Work Environ Health 36, 85–95. 10.5271/sjweh.2897.

20. Navoly, G., and McCann, C.J. (2021). Dynamic integration of enteric neural stem cells in ex vivo organotypic colon cultures. Sci Rep 11, 15889. 10.1038/s41598-021-95434-4.

21. Laible, E., Wegner, A., Knutson, K., Kacmaz, H., Garramone, G.K., Gogineni, K., Matveyenko, A., Linden, D.R., Farrugia, G., and Beyder, A. (2024). Circadian rhythm and whole gut transit in mice. Neurogastroenterol Motil 36, e14771. 10.1111/nmo.14771.

22. Bellampalli, S.S., Fonar, G., Grynyshyn, M., Mercado-Perez, A., Muchhala, K.H., Farrugia, G., Matveyenko, A.V., Linden, D.R., and Beyder, A. (2025). Sex-Dependent Circadian Rhythm Impact on Murine Gastrointestinal Transit. Neurogastroenterol Motil, e70114. 10.1111/nmo.70114.

23. Lakso, M., Pichel, J.G., Gorman, J.R., Sauer, B., Okamoto, Y., Lee, E., Alt, F.W., and Westphal, H. (1996). Efficient in vivo manipulation of mouse genomic sequences at the zygote stage. Proc Natl Acad Sci U S A 93, 5860–5865. 10.1073/pnas.93.12.5860.

24. Obata, Y., Castano, A., Fallesen, T.L., Bon-Frauches, A.C., Boeing, S., Huseynova, A., McCallum, S., Lasrado, R., Heanue, T.A., and Pachnis, V. (2022). Molecular profiling of enteric nervous system cell lineages. Nat Protoc 17, 1789–1817. 10.1038/s41596-022-00697-4.

25. Yamazaki, S., and Takahashi, J.S. (2005). Real-time luminescence reporting of circadian gene expression in mammals. Methods Enzymol 393, 288–301. 10.1016/S0076-6879(05)93012-7.

26. Pendergast, J.S., Niswender, K.D., and Yamazaki, S. (2012). Tissue-specific function of Period3 in circadian rhythmicity. PLoS One 7, e30254. 10.1371/journal.pone.0030254.

27. Morarach, K., Mikhailova, A., Knoflach, V., Memic, F., Kumar, R., Li, W., Ernfors, P., and Marklund, U. (2021). Diversification of molecularly defined myenteric neuron classes revealed by single-cell RNA sequencing. Nat Neurosci 24, 34–46. 10.1038/s41593-020-00736-x.

28. Rao, M., Nelms, B.D., Dong, L., Salinas-Rios, V., Rutlin, M., Gershon, M.D., and Corfas, G. (2015). Enteric glia express proteolipid protein 1 and are a transcriptionally unique population of glia in the mammalian nervous system. Glia 63, 2040–2057. 10.1002/glia.22876.

29. Klein, S., Seidler, B., Kettenberger, A., Sibaev, A., Rohn, M., Feil, R., Allescher, H.D., Vanderwinden, J.M., Hofmann, F., Schemann, M., et al. (2013). Interstitial cells of Cajal integrate excitatory and inhibitory neurotransmission with intestinal slow-wave activity. Nat Commun 4, 1630. 10.1038/ncomms2626.

30. Muller, P.A., Koscso, B., Rajani, G.M., Stevanovic, K., Berres, M.L., Hashimoto, D., Mortha, A., Leboeuf, M., Li, X.M., Mucida, D., et al. (2014). Crosstalk between muscularis macrophages and enteric neurons regulates gastrointestinal motility. Cell 158, 300–313. 10.1016/j.cell.2014.04.050.

31. Shi, J., Hua, L., Harmer, D., Li, P., and Ren, G. (2018). Cre Driver Mice Targeting Macrophages. Methods Mol Biol 1784, 263–275. 10.1007/978-1-4939-7837-3_24.

32. Deaton, R.A., Bulut, G., Serbulea, V., Salamon, A., Shankman, L.S., Nguyen, A.T., and Owens, G.K. (2023). A New Autosomal Myh11-CreER(T2) Smooth Muscle Cell Lineage Tracing and Gene Knockout Mouse Model-Brief Report. Arterioscler Thromb Vasc Biol 43, 203–211. 10.1161/ATVBAHA.122.318160.

33. Obata, Y., Castaño, Á., Boeing, S., Bon-Frauches, A.C., Fung, C., Fallesen, T., De Agüero, M.G., Yilmaz, B., Lopes, R., Huseynova, A., et al. (2020). Neuronal programming by microbiota regulates intestinal physiology. Nature 578, 284–289. 10.1038/s41586-020-1975-8.

34. Mo, A., Mukamel, E.A., Davis, F.P., Luo, C., Henry, G.L., Picard, S., Urich, M.A., Nery, J.R., Sejnowski, T.J., Lister, R., et al. (2015). Epigenomic Signatures of Neuronal Diversity in the Mammalian Brain. Neuron 86, 1369–1384. 10.1016/j.neuron.2015.05.018.

35. Stokkan, K.A., Yamazaki, S., Tei, H., Sakaki, Y., and Menaker, M. (2001). Entrainment of the circadian clock in the liver by feeding. Science 291, 490–493. 10.1126/science.291.5503.490.

36. Davidson, A.J., Poole, A.S., Yamazaki, S., and Menaker, M. (2003). Is the food-entrainable circadian oscillator in the digestive system? Genes Brain Behav 2, 32–39. 10.1034/j.1601-183x.2003.00005.x.

37. Izumo, M., Pejchal, M., Schook, A.C., Lange, R.P., Walisser, J.A., Sato, T.R., Wang, X., Bradfield, C.A., and Takahashi, J.S. (2014). Differential effects of light and feeding on circadian organization of peripheral clocks in a forebrain Bmal1 mutant. Elife 3. 10.7554/eLife.04617.

38. Obata, Y., and Pachnis, V. (2016). The effect of microbiota and the immune system on the development and organization of the enteric nervous system. Gastroenterology 151, 836–844.

39. Macpherson, A.J., Pachnis, V., and Prinz, M. (2023). Boundaries and integration between microbiota, the nervous system, and immunity. Immunity 56, 1712–1726. 10.1016/j.immuni.2023.07.011.

40. Roth, B.L. (2016). DREADDs for Neuroscientists. Neuron 89, 683–694. 10.1016/j.neuron.2016.01.040.

41. Thompson, K.J., Khajehali, E., Bradley, S.J., Navarrete, J.S., Huang, X.P., Slocum, S., Jin, J., Liu, J., Xiong, Y., Olsen, R.H.J., et al. (2018). DREADD Agonist 21 Is an Effective Agonist for Muscarinic-Based DREADDs in Vitro and in Vivo. ACS Pharmacol Transl Sci 1, 61–72. 10.1021/acsptsci.8b00012.

42. Xu, P., Berto, S., Kulkarni, A., Jeong, B., Joseph, C., Cox, K.H., Greenberg, M.E., Kim, T.K., Konopka, G., and Takahashi, J.S. (2021). NPAS4 regulates the transcriptional response of the suprachiasmatic nucleus to light and circadian behavior. Neuron 109, 3268–3282 e3266. 10.1016/j.neuron.2021.07.026.

43. Wen, S., Ma, D., Zhao, M., Xie, L., Wu, Q., Gou, L., Zhu, C., Fan, Y., Wang, H., and Yan, J. (2020). Spatiotemporal single-cell analysis of gene expression in the mouse suprachiasmatic nucleus. Nat Neurosci 23, 456–467. 10.1038/s41593-020-0586-x.

44. Acosta-Rodriguez, V.A., Rijo-Ferreira, F., van Rosmalen, L., Izumo, M., Park, N., Joseph, C., Hepler, C., Thorne, A.K., Stubblefield, J., Bass, J., et al. (2024). Misaligned feeding uncouples daily rhythms within brown adipose tissue and between peripheral clocks. Cell Rep 43, 114523. 10.1016/j.celrep.2024.114523.

45. Wang, H., van Spyk, E., Liu, Q., Geyfman, M., Salmans, M.L., Kumar, V., Ihler, A., Li, N., Takahashi, J.S., and Andersen, B. (2017). Time-Restricted Feeding Shifts the Skin Circadian Clock and Alters UVB-Induced DNA Damage. Cell Rep 20, 1061–1072. 10.1016/j.celrep.2017.07.022.

46. Shan, Y., Abel, J.H., Li, Y., Izumo, M., Cox, K.H., Jeong, B., Yoo, S.H., Olson, D.P., Doyle, F.J., 3rd, and Takahashi, J.S. (2020). Dual-Color Single-Cell Imaging of the Suprachiasmatic Nucleus Reveals a Circadian Role in Network Synchrony. Neuron 108, 164–179 e167. 10.1016/j.neuron.2020.07.012.

47. Smith, C.B., van der Vinne, V., McCartney, E., Stowie, A.C., Leise, T.L., Martin-Burgos, B., Molyneux, P.C., Garbutt, L.A., Brodsky, M.H., Davidson, A.J., et al. (2022). Cell-Type-Specific Circadian Bioluminescence Rhythms in Dbp Reporter Mice. J Biol Rhythms 37, 53–77. 10.1177/07487304211069452.

48. Jacobson, A., Yang, D., Vella, M., and Chiu, I.M. (2021). The intestinal neuro-immune axis: crosstalk between neurons, immune cells, and microbes. Mucosal Immunol 14, 555–565. 10.1038/s41385-020-00368-1.

49. Gabanyi, I., Muller, P.A., Feighery, L., Oliveira, T.Y., Costa-Pinto, F.A., and Mucida, D. (2016). Neuro-immune Interactions Drive Tissue Programming in Intestinal Macrophages. Cell 164, 378–391. 10.1016/j.cell.2015.12.023.

50. Viola, M.F., Chavero-Pieres, M., Modave, E., Delfini, M., Stakenborg, N., Estevez, M.C., Fabre, N., Appeltans, I., Martens, T., Vandereyken, K., et al. (2023). Dedicated macrophages organize and maintain the enteric nervous system. Nature 618, 818–826. 10.1038/s41586-023-06200-7.

51. Mearin, F., Zarate, N., Sardi, J.A., Moreno-Osset, E., and Salis, G. (2003). Traveler’s constipation. Am J Gastroenterol 98, 507–509. 10.1111/j.1572-0241.2003.07249.x.

52. Sack, R.L. (2009). The pathophysiology of jet lag. Travel Med Infect Dis 7, 102–110. 10.1016/j.tmaid.2009.01.006.

53. Hu, L., Li, G., Shu, Y., Hou, X., Yang, L., and Jin, Y. (2022). Circadian dysregulation induces alterations of visceral sensitivity and the gut microbiota in Light/Dark phase shift mice. Front Microbiol 13, 935919. 10.3389/fmicb.2022.935919.

54. Amara, J., Saliba, Y., Hajal, J., Smayra, V., Bakhos, J.J., Sayegh, R., and Fares, N. (2019). Circadian Rhythm Disruption Aggravates DSS-Induced Colitis in Mice with Fecal Calprotectin as a Marker of Colitis Severity. Dig Dis Sci 64, 3122–3133. 10.1007/s10620-019-05675-7.

55. Tran, L., Jochum, S.B., Shaikh, M., Wilber, S., Zhang, L., Hayden, D.M., Forsyth, C.B., Voigt, R.M., Bishehsari, F., Keshavarzian, A., and Swanson, G.R. (2021). Circadian misalignment by environmental light/dark shifting causes circadian disruption in colon. PLoS One 16, e0251604. 10.1371/journal.pone.0251604.

56. Muller, P.A., Matheis, F., Schneeberger, M., Kerner, Z., Jove, V., and Mucida, D. (2020). Microbiota-modulated CART(+) enteric neurons autonomously regulate blood glucose. Science 370, 314–321. 10.1126/science.abd6176.

57. Gammon, S.T., Leevy, W.M., Gross, S., Gokel, G.W., and Piwnica-Worms, D. (2006). Spectral unmixing of multicolored bioluminescence emitted from heterogeneous biological sources. Anal Chem 78, 1520–1527. 10.1021/ac051999h.

58. Pendse, M., De Selle, H., Vo, N., Quinn, G., Dende, C., Li, Y., Salinas, C.N., Srinivasan, T., Propheter, D.C., and Crofts, A.A. (2023). Macrophages regulate gastrointestinal motility through complement component 1q. Elife 12, e78558.

59. Griffiths, J.A., Yoo, B.B., Thuy-Boun, P., Cantu, V.J., Weldon, K.C., Challis, C., Sweredoski, M.J., Chan, K.Y., Thron, T.M., Sharon, G., et al. (2024). Peripheral neuronal activation shapes the microbiome and alters gut physiology. Cell Rep 43, 113953. 10.1016/j.celrep.2024.113953.

