## Supplementary material for "A Novel Cell-Type Specific Circadian Reporter Mouse Reveals Self-Sustained Food Entrainable Nature in Enteric Neurons": Sup table1

**Supplemental Table 1**

| | *Ex vivo* phase of peripheral organ tissue explants (mean ± SD shown, where applicable) | | | | | | |
|---|---|---|---|---|---|---|---|
| | Liver | Adrenal | Esophagus | Kidney | Spleen | Small Intestine | Colon |
| E2a-Cre;Per2CBG-LUC (This study) | 18.59 | 15.81 | 17.85 | 16.71 | 21.69 | 19.1±1.84 | 16.9±1.89 |
| PER2::LUC (Pendergast et al, PlosONE 2012) | 16.1±2.17 | 17.1±1.33 | 19.4±0.71 | 16.40±0.90 | 20.9±0.95 | NA | NA |
| PER2::LUC (Taufique et al, Under Review) | NA | NA | NA | NA | NA | 21.6±3.47 | 19.9±2.01 |
